# A common genomic code for chromatin architecture and recombination landscape

**DOI:** 10.1101/293837

**Authors:** Kamel Jabbari, Johannes Wirtz, Martina Rauscher, Thomas Wiehe

**Affiliations:** Institute for Genetics, Biocenter Cologne, University of Cologne, Zülpicher Straße 47a, 50674 Köln, Germany

**Keywords:** Recombination, Linkage Disequilibrium, Meiosis, TADs, Chromatin, Isochores

## Abstract

Recent investigation established a link between DNA sequences and chromatin architecture and explained the evolutionary conservation of TADs (Topologically Associated Domains) and LADs (Lamina Associated Domains) in mammals. This prompted us to analyse the relationship between chromatin architecture and recombination landscapes of human and mouse. The results revealed that: (1) Blocks of elevated linkage disequilibrium tend to coincide with TADs and isochores, indicating co-evolving regulatory elements and genes in insulated neighbourhood; (2) double strand break (DSB) and recombination frequencies increase in GC-rich TADs, whereas recombination cold spots are typical of LADs; (3) binding and loading of proteins which are critical for DSB and meiotic recombination (Spo11, DMC1, H3K4me3 and PRMD9) are higher in GC-rich TADs. One explanation for these observations is that the occurrence of DSB and recombination in meiotic cells are associated to compositional and epigenetic features (genomic code) that are similar to those guiding the architecture of chromosomes in the interphase nucleus of pre-leptotene spermatocytes.

## INTRODUCTION

A relationship between genome organization and recombination was discovered from banding of human chromosomes several decades ago. It led to the observation that R bands and G/R borders were preferred sites of DNA exchanges and the “hot spots” of mitotic chiasmata [1,2]. These observations suggested a correlation with compositional discontinuities (change of GC% along chromosomes) [3-5], which was later extended and validated statistically [6]. It was proposed that transcriptionally active chromosomal domains could be more accessible to the base-mismatch repair complex, studied by Brown and Jiricny [7,8], which preferentially changes mismatched A’s and T’s into G’s and C’s. Such biased mismatch repair, acting during recombination and recognized as gene conversion [9], was posited to be the most likely explanation [5,11] for local GC variation along chromosomes. With accumulating genomic and genetic data, local rates of recombination were later shown to be positively correlated with GC% in human and mouse [12-15].

At a broad scale, recombination is favoured or suppressed in specific mega-base sized regions [16] where linkage disequilibrium (LD) and GC% are dependent on each other, strong LD was found to be typical of GC-poor regions [17]. Moreover, increasing sequence diversity and GC% were shown to be correlated [18]. Whether the correlation between GC level and recombination is driven by GC-biased gene conversion (BGC) [19-21] or local GC-content itself is driving recombination [22-24], is still a matter of debate. In human, hotspots of recombination are associated with local GC spikes (1 to 2 kb in size) and with an increase of the local mutation rate resulting in G or C nucleotides but have no effect on substitution rates [25], consistent with the observation that while local structure of recombination is fast [26,27], rates over broad scales appear to be constrained [28]. In the context of the present study (and whenever appropriate) we distinguish between local, *i.e.* range of kilobases (*e.g.* 1 to 2 kb for BGC), and regional *i.e*. hundreds of kilobases up to megabases scale (*e.g* chromatin loops and isochores).

At such large scales, it was shown that meiotic chromosomal protein loading is modulated by isochores and that R-bands differentially favor double-strand break (DSB) formation during meiosis [29,30]. Along with meiotic DSB repair, the search for homology and the catalysis of strand exchange are likely to be spatially and temporally coordinated to allow for the formation of higher order chromosome structures (for example, synaptonemal complex, chromosome axes and chromatin loops); this is required for the recombination event to successfully join homologous chromosomal regions during leptotene [30-32].

The entry of pre-leptotene spermatocytes into meiosis (leptotene phase) is accompanied by several epigenetic changes; in primates as well as in mice the location of most DSBs is correlated with the trimethylation of histone 3 lysine 4 (H3K4me3) by the DNA binding enzyme PRDM9 [33-37]. Interestingly, in dogs [38] and several fish species, meiotic recombination tends to occur near promoter-like features [39-41] and PRDM9 protein (absent or truncated in several fish species) does not direct recombination events.

A comprehensive large scale view on chromatin landscape and recombination hot-spots can be obtained by aligning chromosome folding maps with recombination frequency profiles. We study the link between recombination and chromatin architecture, in particular the regional scale of Topologically Associated Domains (TADs) [42,43], an assembly of chromatin loops (also called contact domains) and boundaries 0.2-2 Mb in size [42,44-46], which can be resolved into contact domains, 185 Kb in median size [45]. Another procedure, DamID technology, has shown that lamina-associated domains (LADs) have a median size of 500 Kb, are AT-rich and scattered over the whole genome [44,46, 47]. TADs tend to be conserved among tissues; for instance, 65% to 72% of TAD boundaries are common between human embryonic stem cells and IMR90 fibroblasts [42].

In the present work, we integrate data from different resources (DSBs, recombination, LD-blocks, protein binding of and modification, isochores, TADs and LADs) by analysing their distributions along mouse and human chromosomes, to generate a unifying view on chromatin architecture in meiotic and mitotic cells.

## MATERIAL AND METHODS

### Recombination and LD-blocks data sets

Multiparent populations such as the Diversity Outbred (DO) mouse stock represent a valuable resource of large numbers of crossover events accumulated in present day siblings from founder haplotypes. Coordinates and haplotypes of distinct COs and locations of recombination cold spots are obtained from [40]. This study provides a high-density genotype data set from 6886 DO mice spanning 16 breeding generations, in which 2.2 million CO events were localized in intervals with a median size of 28 kb. By filtering out COs identical by descent, the authors obtained a set of 749.560 distinct COs. They used a measure called “CO information score”, that calculate deviation from the expected frequency of COs with respect to founder strain pairs; this score takes larger values at recombination cold spots.

On the other hand, linkage disequilibrium in human populations is measured in terms of the squared correlation r^2^ between the alleles at a pair of loci (*e.g*., [48]). For our analysis, we used the LD-blocks determined by [49] for populations of European (CEU, TSI, GBR, FIN and IBS), African (YRI, LWK and ASW) and East Asian (CHB, JPT and CHS) descent from the 1k human genomes project, phase I [50]. The authors identified 2605, 1467 and 1725 LD-blocks in African, East Asian and European populations, respectively. In addition to LD-blocks, we used the human sex-averaged recombination map available at the ftp site of the 1k genomes project (HapmapII_GRCh37_RecombinationHotspots.tar.gz).

### Isochores and TAD data sets

To study the correlation between chromatin structure and recombination, we obtained TAD boundaries identified by Hi-C experiments on human and mouse embryonic stem cells [42]. We used the UCSC batch coordinate conversion (liftOver at http://genome.ucsc.edu/cgibin/hgLiftOver) to convert isochore coordinates reported by [50,51] from hg18 to hg19 and mouse genome assembly mm9 to mm10 for compatibility with recombination hot/cold spot coordinates reported in [40]. Isochore maps were visualized with draw-chromosome-gc.pl [52] and Hi-C maps using Juicebox [53] and the 3D genome Browser for comparative Hi-C plots [54]. We also used the recently published, high resolution Hi-C data from mouse sperm [54], GEO accession: GSE79230 and mouse embryonic stem cells [55], GEO accession: GSE96107.

### ChIP-Seq and sequencing data

Meiotic DSBs are induced by dimers of the conserved topoisomerase-like protein Spo11 via a transesterase reaction that links a Spo11 molecule to each 5’ end of the broken DNA and release of covalently bound Spo11 to short oligonucleotides (Spo11 oligos) [56]. By sequencing DNA fragments that remain attached to SPO11 and mapping Spo11 oligos reads [57] of mouse spermatocytes, a total of 13.960 DSB hotspots were defined [58]. We used these data to measure frequency distributions of DSBs along chromosomes.

The genomic coordinates of recombination machinery components, namely, DMC1-bound single strand DNA, H3K4me3 and the DNA-binding sites of the zinc finger, histone methyltransferase PRDM9 were obtained from [59] and [60].

### Statistical analysis

We investigated whether TADs, isochores and LD-blocks overlap more than expected by chance. As a quantitative assessment, we performed an association analysis of genomic regions based on permutation tests using the R/Bioconductor package *regioneR* [61]. When performing an association analysis, it is possible to detect associations that do not reflect boundary proximity. With the “local z-score” function [61] one can test whether or not the association between TADs or isochores and LD-blocks is specifically due to the common boundary positions of the analyzed regions. To this end, we performed a permutation test with *regioneR* and computed *p*-values and z-scores iteratively with shifted positions, 500 kb in 5’ and the 3’ direction with respect to a reference position. Plotting average z-scores versus shifted positions, one can visualize how the value of the z-score changes when moving away from the focal site: a peak at the centre indicates that the association is dependent on the specific genomic coordinates while a flat profile indicates diffuse association (for a recent application, see [47]).

## RESULTS

### Association of recombination rate with TADs and isochores

To test for a potential relationship between TADs and the recombination map in mice, we used the Hi-C map constructed from round spermatids [54] and a map of recombination cold spots constructed from a DO mice cohort [40]. Figure 1A shows a remarkable overlap of recombination cold spots, low GC-isochores and chromatin domains from mouse chromosome 17. GC-poor chromosomal domains, which are frequently attached to the interphase lamina (LADs) [44,46,47] coincide with regions of reduced recombination. This is further confirmed by the distance between consecutive DSBs being larger in GC-poor TADs (Figure 1A). This observation is not limited to mouse chromosome 17, a histogram of GC% of cold spots domains from all chromosomes (Figure 1B) shows that the underrepresentation of cold spots in GC-rich TADs is a genome wide feature (mean difference t-test; *p*-value < 2.2e-16). Similar results are obtained with embryonic stem cells (S1 Figure) [55], see also [54]).

**Figure 1.**
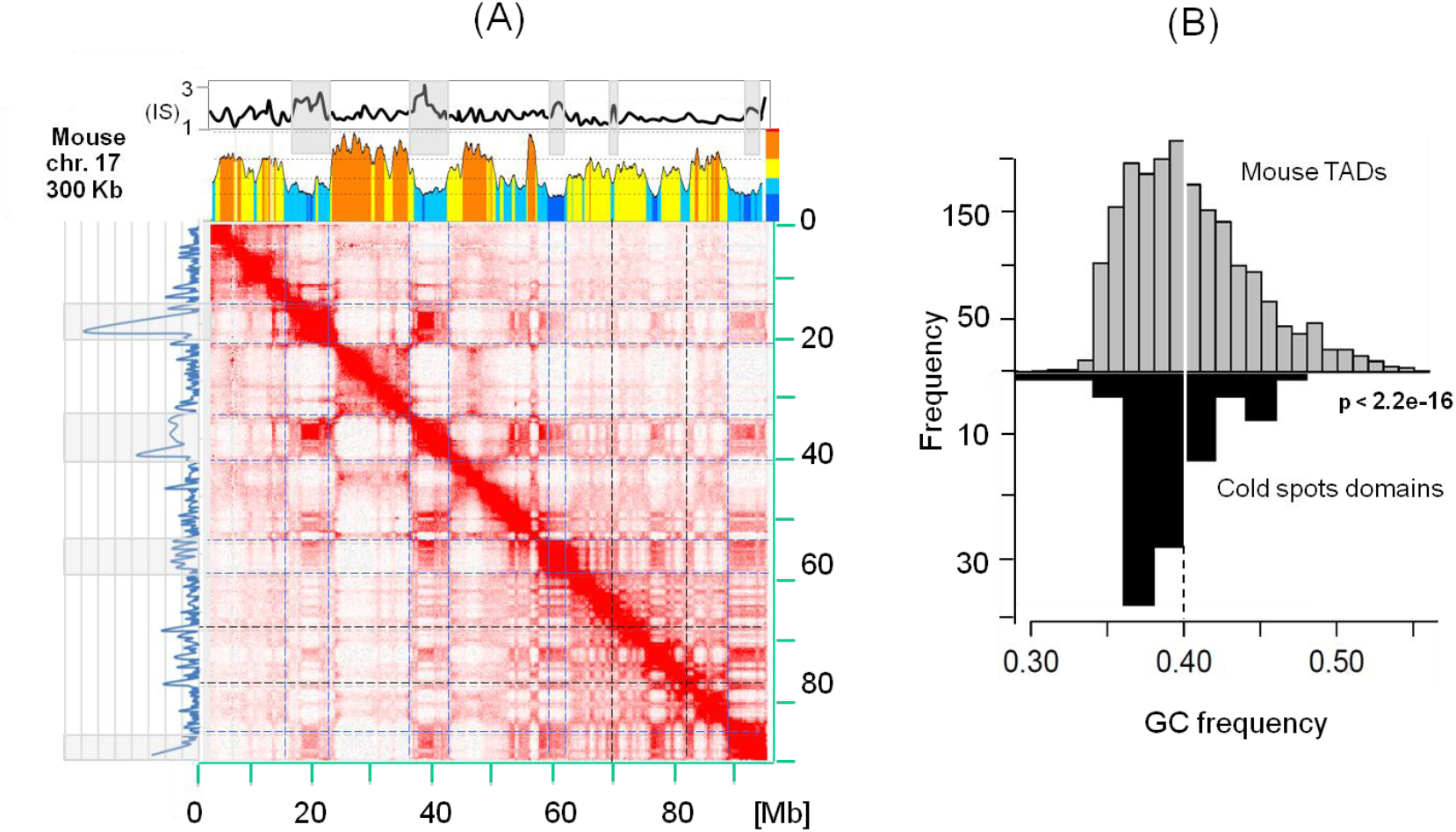
Spatial distribution of recombination cold spots TADs/LADs and isochores. (A) The heat-map of chromatin interactions in mouse chromosome 17 from round spermatids aligned to the profile of cold spot information score (IS) from [40] (top blue curve with grey banded cold spots) and the corresponding compositional profile (the multi-coloured profile). The left vertical panel represents the distance between adjacent spo11 DSB events; large distances are highlighted with grey bands (vertical lines are 500 kb spaced). GC profile is drawn from mm10 genome assembly with a sliding window of 300 Kb (with steps 1/10 of the window size). Increasing GC isochores are represented by deep blue (L1), light blue (L2), yellow (H1) and orange (H2). Blue broken lines across the heat-map delimit cold spots (potential LADs). Black broken line point to distantly positioned DSBs visibly coinciding with abrupt GC drops. (B) GC% of ES cells TADs from [42] (top) and of cold spots intervals (bottom) defined in [40].

In agreement with the results of Figure 1A and 1B, we also observe a significant positive correlation between recombination rate and TAD GC% in mouse (Figure 2A); and human (Figure 2B).

**Figure 2.**
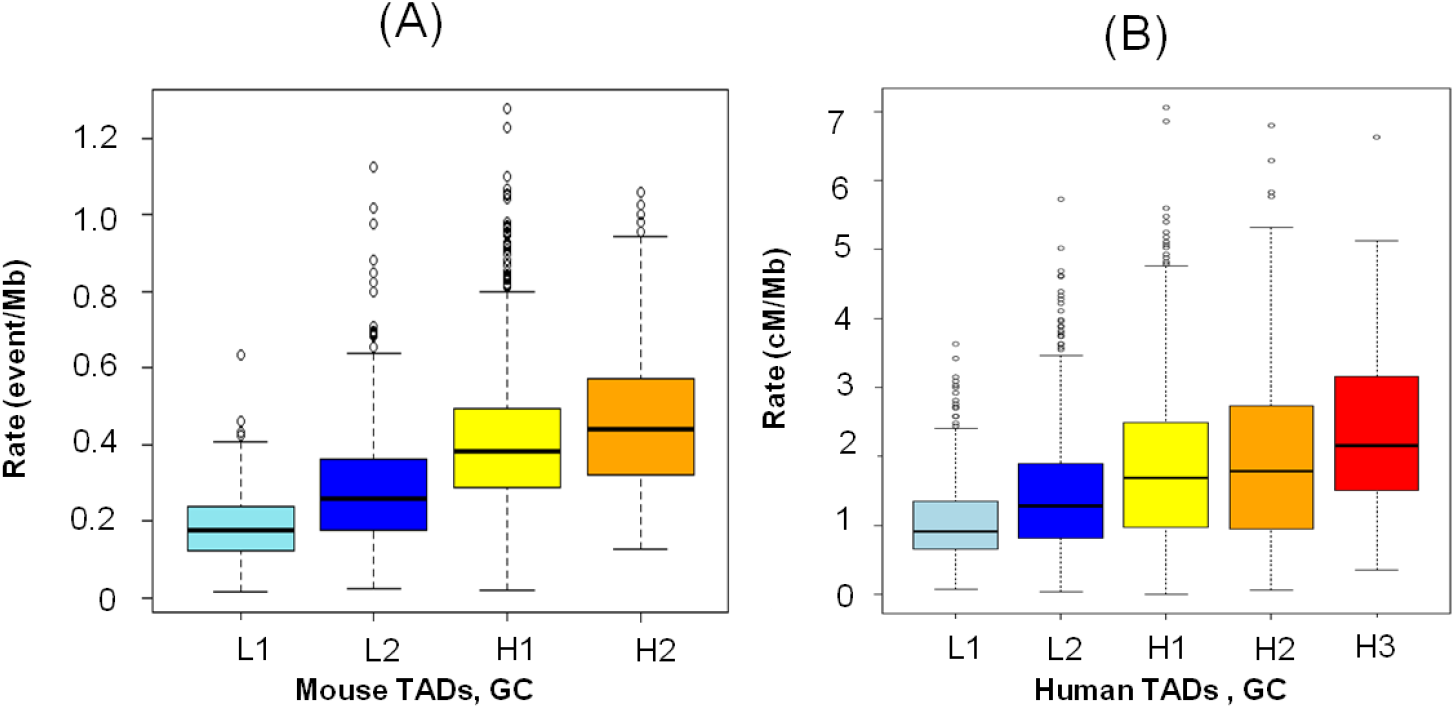
Correlation between recombination rate and isochore or GC level of TADs. (A) Box plot representing the association between TADs GC levels in mouse embryonic stem cells and recombination rate based on data from Morgan et *al.* 2017 [40]. (B) Box plot representing GC increase of human TADs with recombination rate in human. L1, L2, H1, H2 and H3 correspond to human TADs or isochore families. Notice that H3 Isochores are absent in mouse (for more details [51] and [47]).

### Linkage disequilibrium blocks match TADs and LADs

Recombination cold spot intervals and TADs appear to coincide, as shown in the coarse-scale heat-map of Figure 1A. This suggests that regions of low recombination may have elevated levels of linkage disequilibrium. To test this in human, we evaluated the statistical significance of overlap between blocks of linkage disequilibrium (LD), TADs, and isochores. The genome wide statistical test for overlap between TADs or isochores and LD-blocks are all highly significant (S2 Figure), all *p*-values are less than 0.001. To assess the strength of boundary sharing between LD-blocks and TADs, we first established a reference by calculating the z-score profile of shared LD-blocks between European *and* African population. The z-score profiles for shared TADs and LD-blocks boundaries are plotted with respect to this reference (Figure 3).

**Figure 3.**
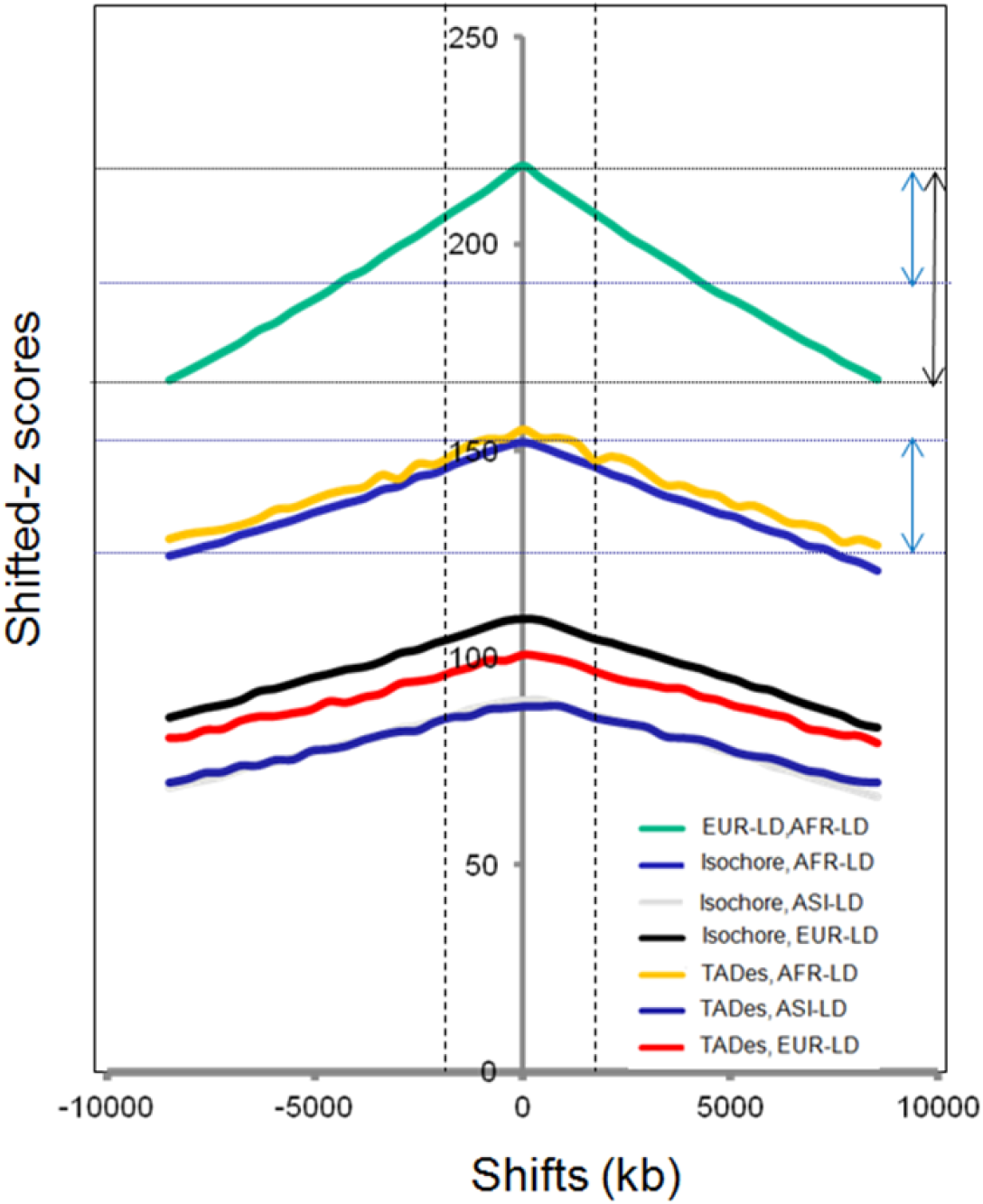
TADs, Isochores and LD-blocks boundaries concordance. Shifted z-score (on the y-axis) changes when moving the region set interval (x-axis): the peak at the centre indicates that the association is dependent on the genomic coordinates, while a flat profile will indicate that is the association either absent or boundary independent. Abbreviations: ASI*-*LD, AFR*-*LD and EUR*-*LD correspond to Asian, African and European linkage disequilibrium blocks, respectively. Black and blue arrows show the difference in drop of z-scores between EUR-LD *vs*. AFR-LD and isochores or TADs *vs*. AFR*-*LD, pointing to a weaker drop of z-scores in the latter compared to the former. The arrows indicate ∼50% drop in z-score compared to the reference (EUR-LD, AFR-LD).

The z-score profiles (Figure 3) clearly indicate a concordance between TADs or isochores and LD-blocks, although weaker in strength (∼50% drop in z-score compared to the reference, see arrows in figure 3). Interestingly, the peak of z-score profile is most pronounced for the ancestral African population (see Discussion).

The regional co-variance of TADs, isochores, recombination or LD blocks can be visualized using an LD heat-map based on a measure called topological LD (tLD); tLD is based on genealogical rather than allelic clustering (see [62] for more details).

As expected the two LD heat maps are very similar (Figure 4A) and the linkage signal is magnified with tLD, which is practical for visual inspection. TADs and LD-blocks boundaries match each other and the two match the GC-poor isochore boundary, probably constrained by early anchoring to the interphase lamina (Figure 4B and S3 Figure). Recombination frequency is also compartmentalized, the GC-rich loop has clearly higher recombination frequency, hence fitting the expectation from the GC *vs*. Recombination positive correlation of Figure 1 and Figure 2. It is important to notice that the loop boundary is conserved across seven cell types analyzed (S3 Figure). The boundary is invariably located in the R3HDM1 gene; the LCT and MCM6 genes are within the short GC-rich TAD.

**Figure 4.**
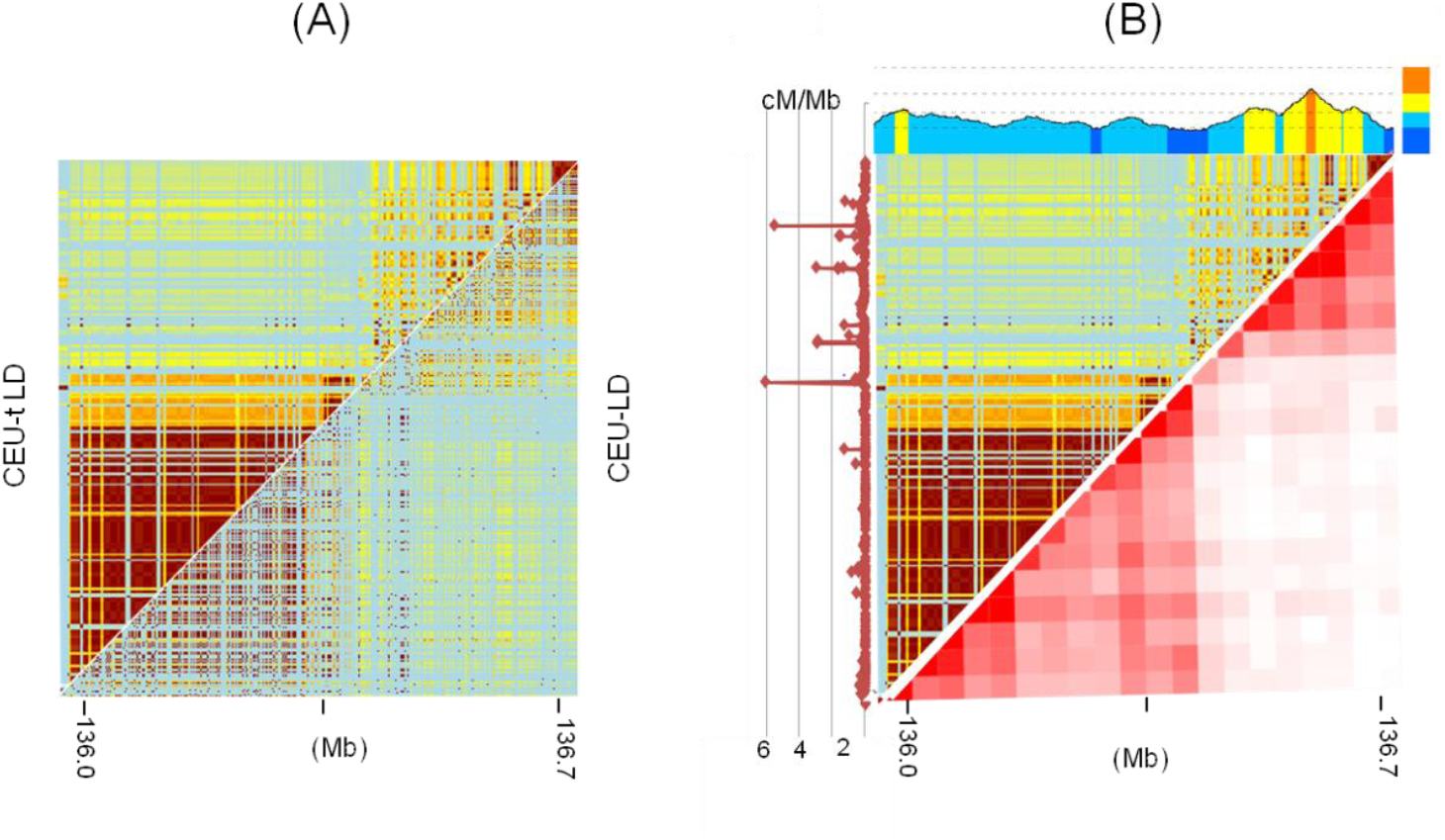
**(A)** The heat maps of conventional (right) and topological LD generated for the LCT and neighbouring regions (chr2:135.9-136.7 Mb) from the CEU subsamples of the human 1k genomes project **(B)** The aligned heat maps of TADs and tLD from a selected region of chromosome 2, encompassing the LCT gene and neighbouring genes of the CEU (top). Left graph (Brown line) represents recombination frequency in cM/Mb.

### Recombination machinery components and TADs

Based on the human and mouse analysis, we hypothesize that the correlation between recombination rate, chromatin loops and GC% (Figures 1 and 2) is a general property of mammalian genomes. In the case of mouse, binding of key recombination proteins mapped around DSB sites [59,60]are also positively correlated with recombination rate and with GC level of TADs (Table 1), moderate to strong positive correlations are observed. All correlation reported in Table 1 are statistically significant and can be considered particularly robust because they include different PRDM9 alleles and different mice strains, and therefore diverse recombination hotspot positions.

**Table 1:** Pearson Correlation coefficients between genomic features. “R” refers to recombination events per 100 kb; numbers in blue and black refer to RJ2 and B6 mouse strains, respectively (data from [60]); numbers in red refer to data from [58] on B6 mouse strains. In all cases *p*-values are less than. 2e-16.

|  | R | Spo11 | Prdm9 | H3K4me3 | DMC1 |
| --- | --- | --- | --- | --- | --- |
| TADs GC% | 0.46 | 0.38 | 0.25-0.32 | 0.35-0.43 | 0.26-0.20 |
| R |  | 0.38 | 0.34-0.39 | 0.22-0.38 | 0.33 |

## DISCUSSION

### Interphase-leptotene chromatin reorganization

In mice and humans, spermatocytes begin to enter leptotene, the first stage of meiotic prophase and later in this stage chromosomes are re-organized in alternating domains of greater and lower DSB activity [63,64]. Leptotene chromatin domains are usually visualized as linear arrays of chromatin loops, connected by the synaptonemal axis (see for example [65,66]), quite different from the interphase chromatin, where about 40% of the genome is associated with the nuclear envelope [44]. Cohesin has an architectural role in the organization of interphase chromosomes and similar roles have been proposed for cohesin and the related condensin complexes in meiotic and mitotic chromosomes [66,67]. This begs an obvious question: Can the pre-existing higher order chromatin structure in the interphase of progenitor germ cells (PGC) or pre-leptotene spermatocytes (PLS) be related to those of early leptotene? The spatial correlation between the recombination rate profile and the loop structure of spermatids and ES cells appears to argue in favour of this link. Indeed, shorter loops are biased towards GC-rich TADs (see Figure 1A), which is consistent with the smaller size of the loops attached to the SC in the telomeric and subtelomeric regions of mouse [68], known to be GC and gene-rich [69-71]. As expected from the lower density of DSBs in LADs, the distance between two consecutive DSBs is longer than that of GC-rich TADs (Figure 1A and S4 Figure), reflecting preferred attachment sites of the SC axial elements. A parsimonious explanation could be that the chromatin loops of many leptotene chromosomes are, in part, inherited from the interphase of PGC or PLS through DNA compositional constraints and active (*e.g.* H3K4me3, H3K9ac and H3K4me2) and/or repressive (*e.g.* H3K9me3, H3K27me3, H3K9me2) chromatin marks ([72] for a review). The latter could maintain these regions in a “poised” folding state participating later in the assembly of chromosome axis and loops.

It is believed that the binding of transcription factors and chromatin modifiers influence DSB density [73,74] and that DSBs occur in regions of accessible chromatin (GC-rich) that are present in mitotic as well as in meiotic cells [75]. These observations are in accordance with previous anticipation that meiotic prophase may have evolved directly from the latter stages of the mitotic program [30]. Because GC-poor TADs/LD-blocks are on average longer and gene poor (richer in genes with large introns), one may conclude that the loops emerging from the synaptonemal complex are of different sizes and may be influenced by chromatin packing [76]. DSBs typically occur in loop sequences [77] and their chromosomal distribution could be affected by loop size [60,78] and nucleotide composition of loops, as shown Figure 1.

At shorter scales, nucleosome formation generally restricts the accessibility of proteins to DNA, including Spo11. Meiotic DSBs are reportedly introduced on the chromatin loop regions that transiently interact with the lateral elements of the synaptonemal complex [77,64,79], suggesting that these chromatin regions may contain nucleosome-depleted regions (GC-rich open chromatin) [80-82,60]. Thus, the local recombination effects are expected to be stronger in GC-rich TADs, where loops are shorter [47,83], nucleosomes are spaced, and PRDM9 (when present) and CTCF binding sites are enriched [60]. It should be noticed here that although a functional PRDM9 homolog is reported to be missing in dogs [38], recombination cold spots genomic distribution recapitulates the major features of the mouse recombination map [40], in particular, recombination appears to be directed to gene promoters and CpG islands. This is in agreement with the observation that isochores and TADs are conserved between human, dog and mouse [42,45,84,47] and suggests that the recombination machinery may follow the principles of 3D genome organization in their selection of recombination partners as it does after damage-induced DSBs, which is primarily dictated by genome shape and nuclear position [85,86].

### LD-blocks and chromatin neighbourhoods

The concordance between TADs or isochores and recombination or LD-blocks revealed in this work has two consequences; (1) it suggests that regional variation of recombination is topologically defined in concert with an underlying compositional and epigenetic framework (common genomic code). GC-rich domains are recombinogenic, which may explain the small size of GC-rich LD-blocks and TADs (S5 Figure). As a result, shared LD-blocks between Europeans, East Asians and Africans, are significantly larger in size (S6 Figure) and their lower recombination rate is in agreement with the fact that strong LD blocks are GC-poor [17] and enriched in recombination cold spots as shown in Figure 1; strong LD between a pair of SNPs in GC-rich recombinogenic TADs may hint to functional chromatin contacts maintained by purifying selection. A reduced recombination rate between target genes and their regulatory elements may come with an advantage for genes and regulatory elements under stronger selection in the germ line lineage, possibly by maintaining multiple alleles as a single inheritance unit [87]. The ability of LD-blocks to encompass long range SNP interactions in regions with enhanced intra-loop contacts, highlights how allelic chromatin topology analyses can help to infer mechanisms by which SNPs associate with disease and traits, through haplotypes mapping of chromatin Interactions. On way is to assess disease association of disrupted CTCF-mediated interactions by examining linkage disequilibrium between the SNPs residing in CTCF motifs and GWAS disease-associated SNPs [88].

Chromosomal segments have been broken down by repeated meioses along human evolution, assuming the overall level of LD is closer to recombination-drift equilibrium in the ancestral African population than in the derived European and Asian ones, our observation of a distinct z-score profile in African on one hand and European and Asian on the other, (Figure 3) is consistent with this assumption and agrees with the homozygosity runs (ROH) being longer in European and Asian compared to African [89]. LD-blocks like ROH of much older origin, are generally much shorter and longer GC-poor isochres may persist due to regionally low recombination rates. Yet, the exact nature and molecular basis of this genetic and structural (TADs and LADs) link remain open to investigation.

### Chromatin organization imposes structural constraints on recombination

The majority of recombination cold spots are found in pre-leptotene constitutive LADs (the GC-poorest isochores), which are enriched for structural variations, in particular deletions [90,40], that can locally suppress recombination due to miss-alignments of homologous chromosomes disrupting synapsis [40]. Loops will be stabilized as in mitotic chromosomes, by the dynamic binding of cohesin and meiotic insulators (*e.g*. CTCF) [91,92]. This is in keeping with recent studies showing that BORIS (or CTCF-like), a genomic neighbour of Spo11 (CTCF-like and Spo11 are located in two nested neighbouring loops [93]), is present in male germ cells during and after meiosis, and may interact with at least one of the meiosis-specific subunits of cohesin complexes, hence contributing to a progressive re-establishment of genome architecture in haploid post-meiotic round spermatids [94,54].

The trade-off between major determinants of the genomic code, namely, DNA sequence constraints and its modified or native protein binding partners, is expected to affect the conformation of the chromatin fibre, believed to be primarily determined by its own stiffness, favouring or restricting the formation of long-range contacts [30]. Equally interesting in this regard are the compositional constraints that shape DNA bendability [95] and super-coiling [96].

## CONCLUSIONS

The frequency of hot spots of recombination and its relation to open chromatin was suspected many decades ago. The recent observation of the correspondence between isochores and TADs [47] and the large number of available genomic maps allowed us to investigate and demonstrate that LD-blocks significantly, albeit weakly, match TADs and isochores. Cold/hot spots of recombination are compartmentalized, they correspond to interphase AT-/GC-rich TADs. Binding affinities of key determinants of meiotic recombination hot spots (PRMD9, Spo11, DMC1, H3K4me3) are positively correlated with TADs GC%. This suggests that recombination frequency is associated with the same compositional and epigenetic features that constrain the distribution of chromatin in the interphase nucleus. In sum, these observations indicate that the recombination landscape in mammals and the allele co-segregation associated to it, are tied to insulated interactions in which chromatin folding is crucial. Pre-meiotic TADs and sub-TADs and there sequence design (*e.g*. oligonucleotides frequencies) might represent a structural template within which meiotic loops organize themselves, evoking a “meiotic topological memory”.

## Footnote

We would like to thank Tomaz Berisa for sharing data on linkage disequilibrium blocks in human populations.

Lately LD-blocks and TADs have also been investigated in two other studies which were deposited on bioRxiv. The first study [97] did not find a significant correlation of TAD boundaries with classical LD blocks, but when applying a new measure, called “Linkage Probability”, the authors could report a strong association. The second article [98] did not detect such an association.

## SUPPROTING INFORMATION CAPTIONS

### Supplemental S1-S6 Figures

**Figure S1:**
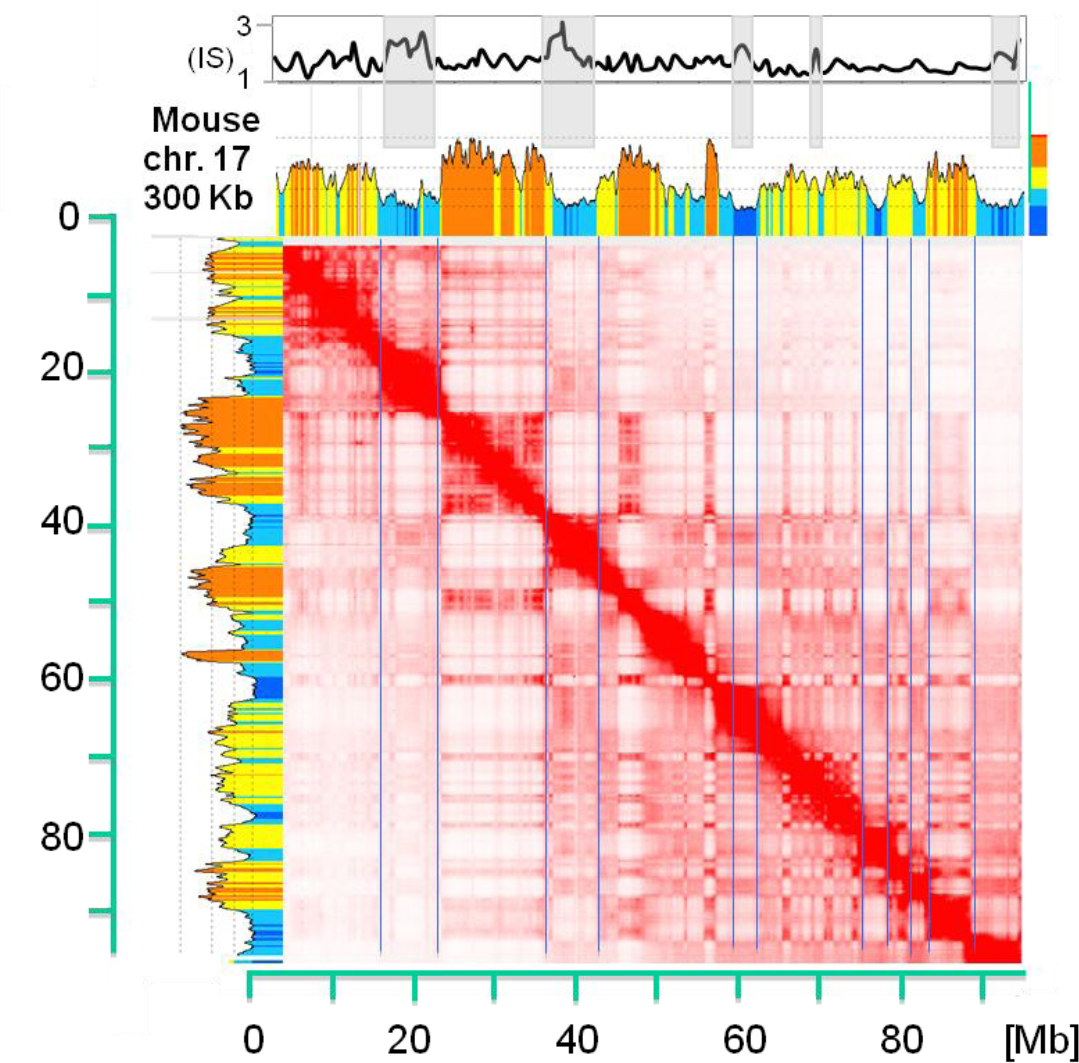
Spatial match between recombination rate, TADs and Isochores. The heat-map of chromatin interactions in mouse embryonic stem cells of chromosome 17 is from Bonev *et al.* 2017. Other annotations are as in Figure 1.

**Figure S2:**
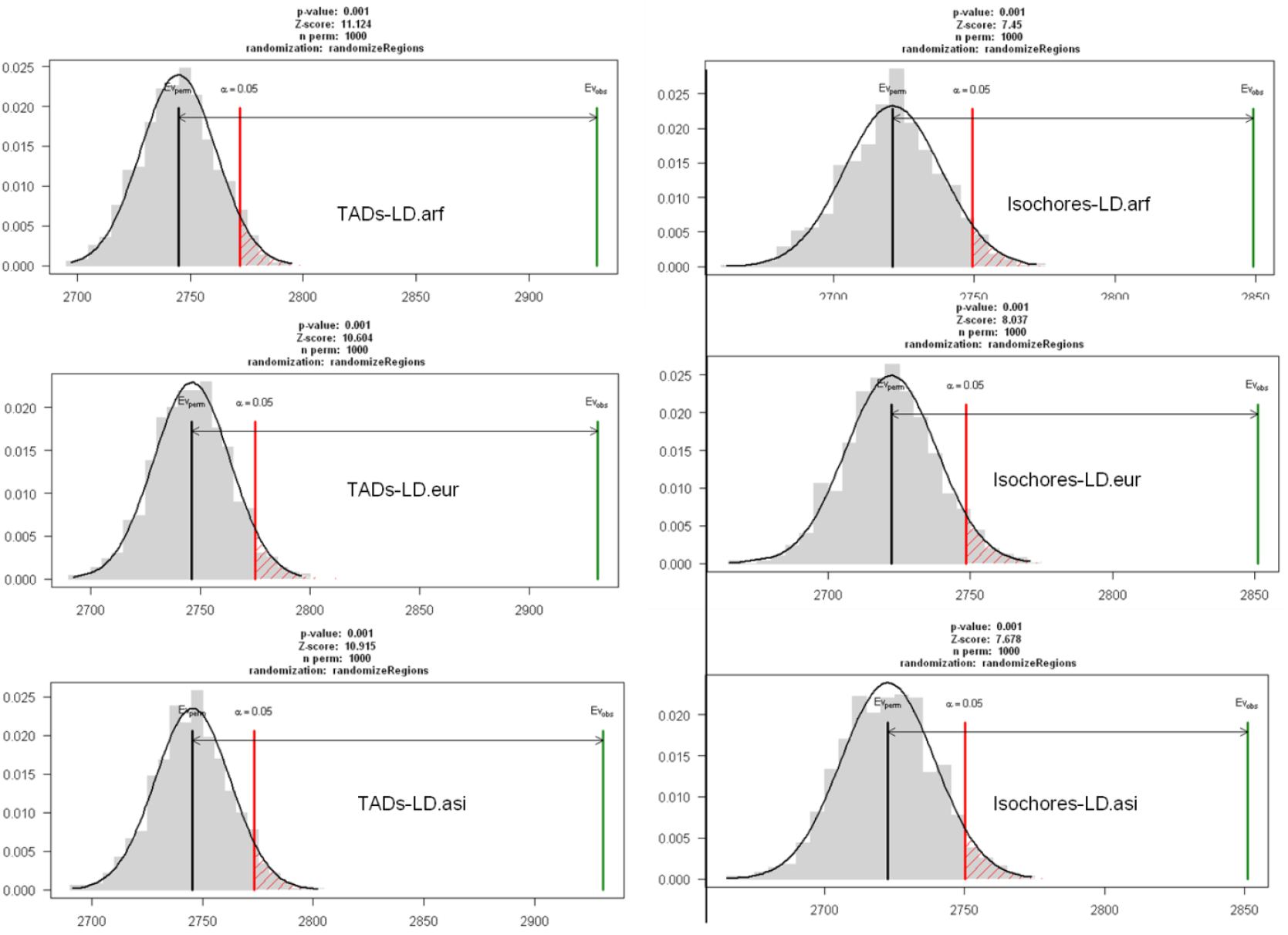
Association analysis of genomic regions based on permutation test. Human LDs from African (afr), European (eur) an East Asian (asi) overlap significantly with TADs from human ESC cells and isochores.

**Figure S3:**
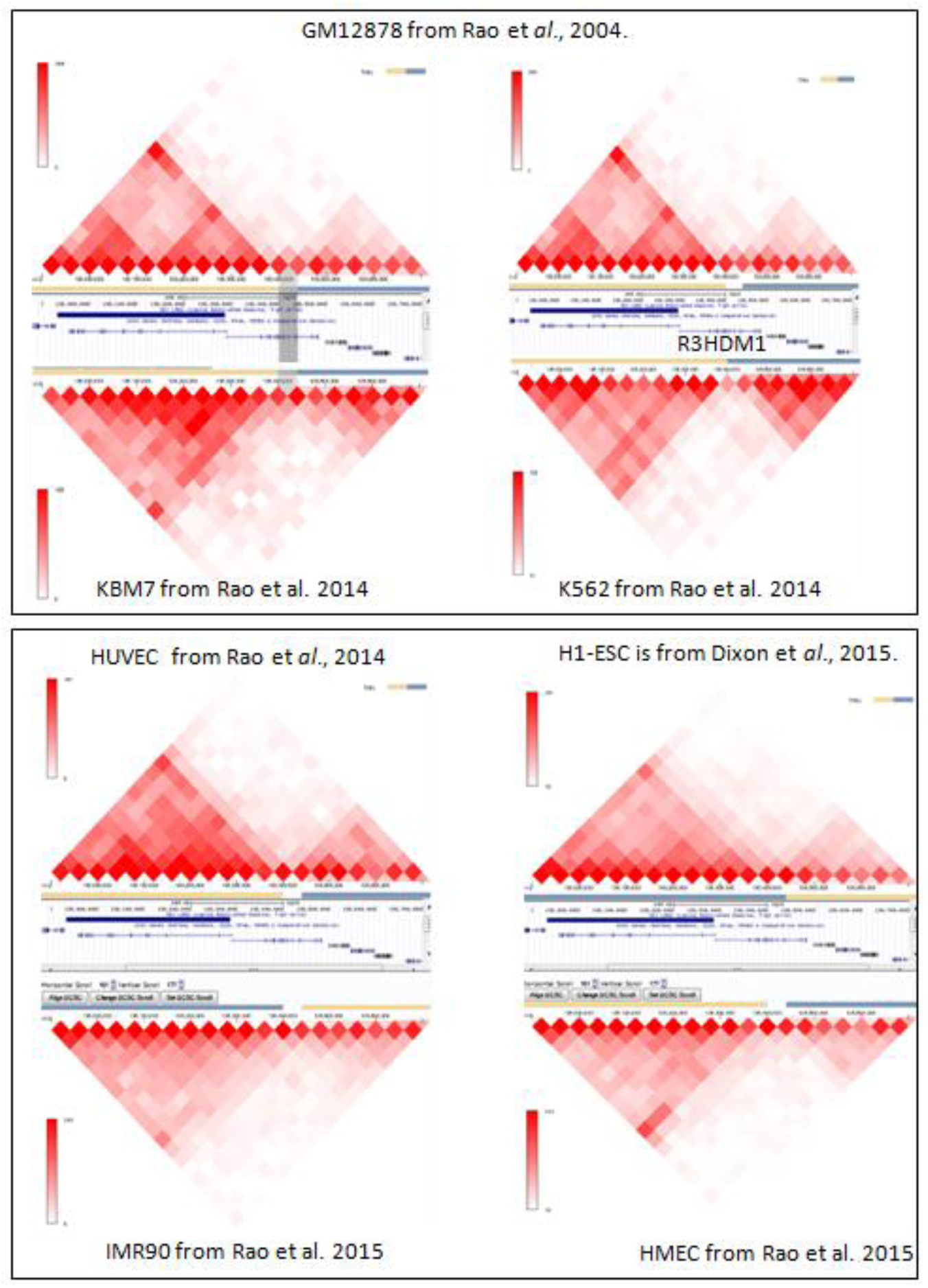
Chromatin loop boundary stability along the LCT locus (chr2:135.912.665-136.712.228). Beige and light blue lines indicate TADs (visualized using the 3D Genome Browser). Similar topology is noticeable with the gene R3HDM1 at the domains boundary. Blue horizontal bar (under the TADs annotation bars) represents annotated LAD from TIG-3 (RRID:CVCL_E939) cell lines.

**Figure S4:**
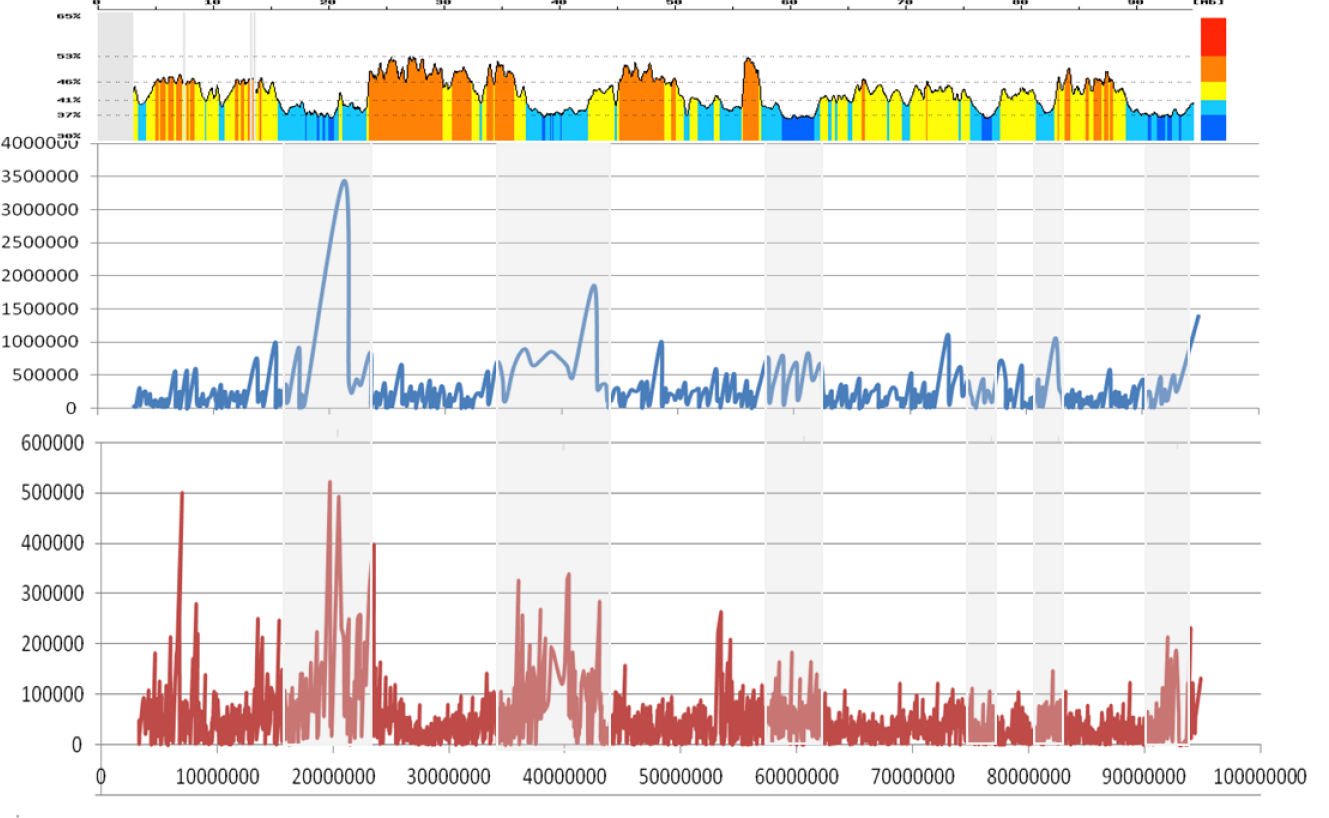
Distance between adjacent spo11 DSB events (blue) and adjacent recombination events (brown) of mouse chromosome 17; grey bands correspond to cold spots dense areas. Top panel is the isochore map; other indications as in Figure 1.

**Figure S5:**
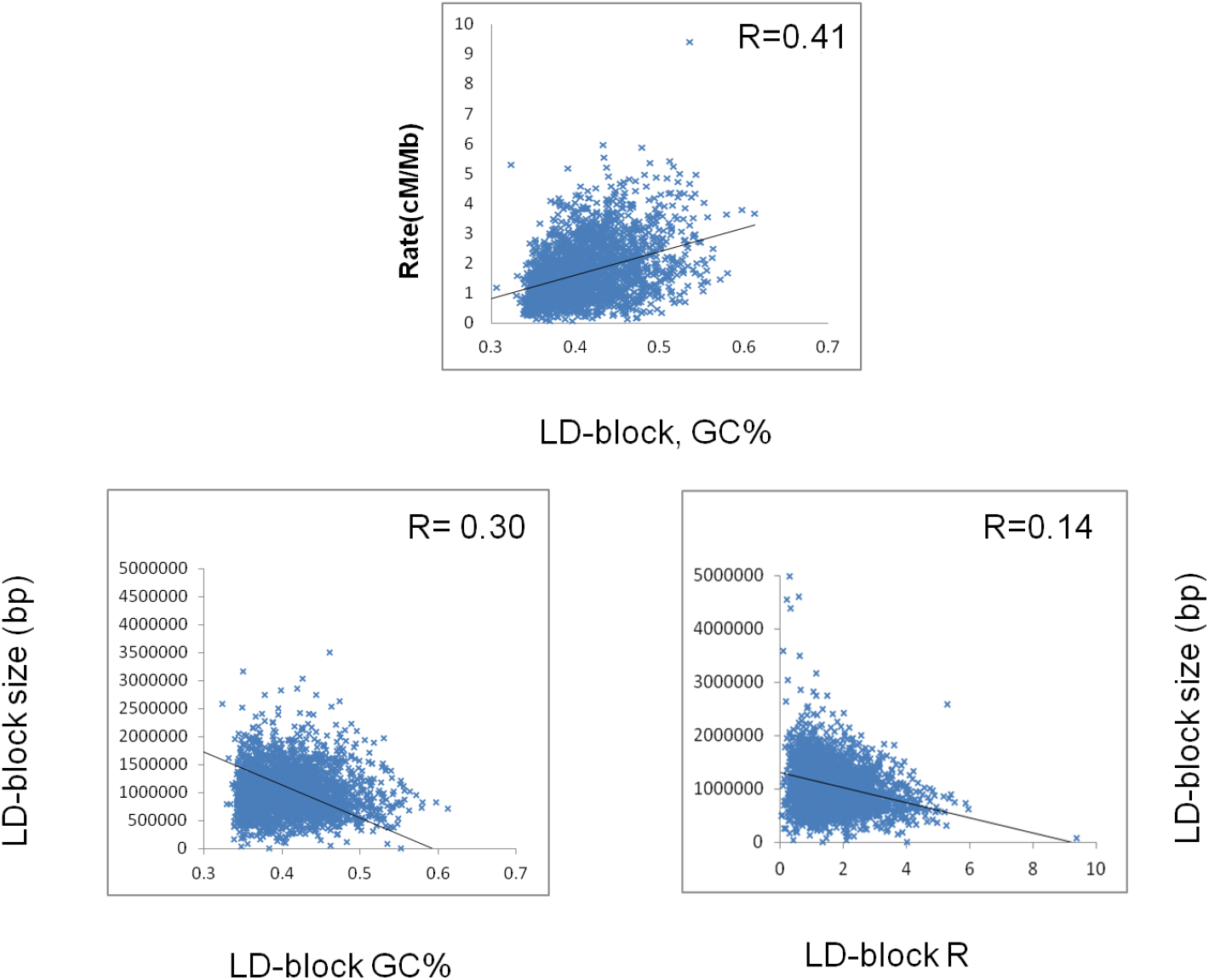
Correlation between LD-blocks GC%, LD-block size and recombination rate.

**Figure S6:**
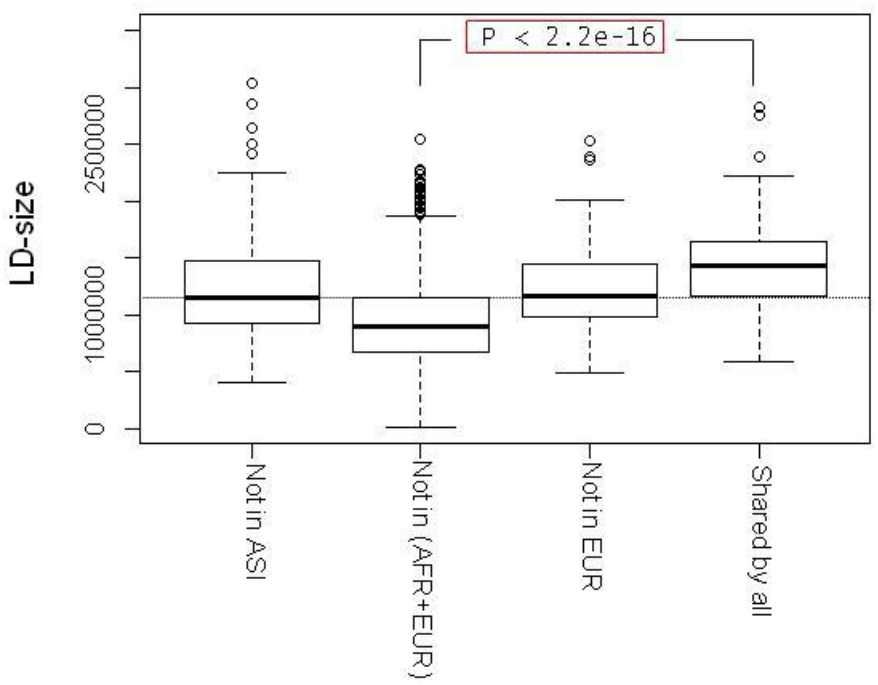
Box plot showing significant (p-value < 2. 2e-16) increase of shared LD-blocks size; median=1. 3 Mp for “Shared by all” LDs *vs.* 0.9 Mb for “Not in (EUR+AFR)”.

